# In-depth 3D Exploration of Autosomal Dominant Polycystic Kidney Disease Through Light Sheet Fluorescence Microscopy

**DOI:** 10.1101/2025.03.18.644002

**Authors:** Pablo Delgado-Rodriguez, Itsaso Vitoria, Gonzalo R. Ríos-Muñoz, Lídia Bardia, Nicolás Lamanna-Rama, Laura Nicolas-Saenz, Jon Sporring, María L. Soto-Montenegro, Rafael Aldabe, Julien Colombelli, Arrate Muñoz-Barrutia

## Abstract

Autosomal Dominant Polycystic Kidney Disease (ADPKD) is the most prevalent genetic kidney disorder. Animal preclinical studies are one of the main tools to study this disease, often through either 2D histology imaging for high-resolution analysis or CT or MRI for full kidney segmentation. As an alternative to these modalities, we propose the use of Light Sheet Fluorescence Microscopy (LSFM) for high-resolution 3D imaging of healthy and ADPKD-induced mouse kidneys, enabling a detailed volumetric morphological analysis of the disease’s effects. In a mouse ADPKD model, *ex vivo* imaging of the kidneys was performed through LSFM, after which a combination of machine learning and other processing techniques allowed us to perform an in-depth image analysis. This includes the segmentation of key structures, such as the full kidney volume and, within it, its internal cavities, cortex, glomeruli, and cysts, complemented by texture analysis of tubular structures in the cortical area. Pathological kidneys exhibited significant volume enlargement and increased internal cavities due to cystogenesis. While glomerular count remained stable, their spatial distribution was altered, showing increased interglomerular distances and showcasing the deformations produced by the disease. The texture analysis of tubules from the cortex region identified Local Binary Pattern (LBP) uniformity and porosity as key biomarkers of tissue deformation, which could be used as markers to further evaluate the development of the disease. These findings underscore the potential of LSFM imaging as a powerful tool for detailed ADPKD characterization and treatment assessment.

## 1 Introduction

Autosomal Dominant Polycystic Kidney Disease (ADPKD) is the most common kidney hereditary disease, characterized by the development of cysts that progressively deform and enlarge the kidney, impairing its normal function [1]. It is produced by the mutation of either the Polycystic Kidney Disease gene 1 (PKD1) or the Polycystic Kidney Disease gene 2 (PKD2). Current diagnosis in humans is initially performed on patients with a known ADPKD family history [2]. Ultrasound (US) imaging is the first tool to be applied, as a non-invasive and inexpensive technique, with ADPKD being diagnosed depending on the amount of cysts found per kidney. For non-conclusive cases, Magnetic Resonance Imaging (MRI) is employed as a more precise tool that also allows the observation of cysts that could have been missed through the US alone. There is no current cure for this condition, with treatments focusing on slowing down cyst formation, mainly through the use of Tolvaptan, somatostatin analogs, or more experimental treatments like Sodium-glucose cotransporter inhibitors (SGLTi) [3].

Imaging plays a crucial role in ADPKD research [4], with disease progression primarily assessed through Total Kidney Volume (TKV) measurements. MRI is the preferred modality for this purpose [5], although Computed Tomography (CT) serves as an alternative, particularly when MRI is contraindicated. Additionally, 3D ultrasound has demonstrated the ability to provide comparable TKV measurements [6]. In recent years, deep learning techniques have been widely applied to TKV quantification [7], often in combination with additional features and biomarkers to enhance disease prognosis [8]. Furthermore, deep learning-based cyst segmentation has been applied to clinical MRI [9, 10, 11] or CT images [12], offering new tools for assessing ADPKD progression.

Despite these advances, clinical studies on ADPKD face practical and ethical constraints that limit the exploration of underlying disease mechanisms and the evaluation of new treatments. This underscores the need for preclinical animal studies, typically conducted on mice or rats under controlled conditions. In these studies, histology imaging has provided high-resolution insights into cyst formation, enabling investigations into disease degradation mechanisms [13], the effects of novel drugs [14, 15], and genetic modifications [16, 17], as well as facilitating disease characterization through temporal imaging modalities [18]. Cyst segmentation in 2D histology images has also been used to assess ADPKD progression via the cystic index, which quantifies the proportion of cyst area relative to the total tissue [15, 19], with various computational tools developed for this purpose [20, 21]. However, 2D histological imaging provides only a limited approximation, as it captures information from specific slices of the organ. In contrast, three-dimensional modalities such as CT [22] and MRI [23] enable comprehensive monitoring of total kidney volume and disease burden, offering a more complete view of disease progression.

To characterize and evaluate volumetric changes in ADPK-affected kidney tissue at a finer scale, combining high resolution with whole-organ 3D reconstruction, we propose an alternative imaging approach: Light Sheet Fluorescence Microscopy (LSFM) [24] applied to optically cleared kidneys. LSFM is a fluorescence-based microscopy technique in which a laser beam is shaped into a planar illumination. This sheet selectively illuminates a specific plane of the sample, exciting fluorescent molecules that emit light captured by the camera positioned at a 90º angle to the light source. By translating the sample across the light sheet, LSFM enables the generation of a high-resolution 3D reconstruction. For optimal imaging, the sample must be as transparent as possible to minimize both excitation and emitted light scattering (for a review, [25]). When tissues are not naturally transparent, a clearing process is applied, involving the removal of proteins and/or lipids and their replacement with substances of matching refractive index to reduce scattering, thereby enabling one to image across whole organs and organisms [26]. In this study, we aim to observe and quantify the effects of ADPKD on mouse kidneys using LSFM, requiring their extraction and preparation into optically cleared *ex vivo* samples [27]. LSFM can generate highly detailed 3D reconstructions and has been widely applied to study whole organs and tissues such as the brain [28, 29, 30]. However, its application to kidney imaging remains scarce, despite promising results demonstrating its potential for capturing high-resolution structural data [31, 32].

In this manuscript, we propose the use of LSFM for *ex vivo* 3D imaging of mouse kidneys, providing a detailed preclinical assessment of ADPKD. This approach combines a high optical resolution approach, comparable to histology imaging, but with volumetric capabilities akin to modalities like MRI, to enable a comprehensive three-dimensional analysis of kidney architecture. By integrating advanced computational processing techniques, including machine learning, we aim to extract quantitative insights into the structural effects of ADPKD. Specifically, this study focuses on segmenting and quantifying key kidney components, such as glomeruli, cysts, and internal cavities, while also employing texture analysis of lower-scale tubular regions in the cortex to further characterize disease progression at a microscopic level. These analyses provide a novel framework for evaluating disease severity and therapeutic responses, with the potential to enhance preclinical nephrology research.

## 2 Materials and Methods

### Generation of PKD2 Knockout Mice and Sample Preparation

All experimental animal procedures were conducted according to European Communities Council Directive 2010/63/EU and ARRIVE guidelines [33] and approved by the Ethics Committee for Animal Experimentation of Hospital Gregorio Marañón and University of Navarra.

We used Pax8rtTA [34], TetO-Cre [35], PKD2fl/fl mice, provided by the Baltimore Polycystic Kidney Disease (PKD) Research and Clinical Core Center. A conditional inducible knockout (KO) was produced for the PKD2 gene to induce the disease. The transgenic model contains loxP sites flanking exons 11-13 of the PKD2 gene, which can be targeted by Cre recombinase under the control of a tetracycline-responsive promoter element (tetO).

Upon administration of doxycycline (DCX), a tetracycline analog, the reverse tetracycline-controlled transactivator protein (rtTA) is expressed, triggering Cre-mediated recombination and removal of the floxed PKD2 sequence. This system enables tissue-specific deletion of the PKD2 gene, as the Pax8 promoter drives expression in the proximal and distal kidney tubules, as well as in the collecting ducts of the kidney.

For this study, both healthy and ADPKD-induced mice were bred at Centro de Investigación Médica Aplicada (CIMA) Universidad de Navarra (Spain), and subsequently transported to Hospital General Universitario Gregorio Marañón. When the mice reached 12 weeks without DCX, they were injected with DyLight™ 594 labeled tomato lectin (0.2 µg/µl, DL-1177-1, Vectorlabs, Spain). Tomato lectin is an effective marker of blood vessels and its conjugation with a fluorophore enhances the visualization of intravascular structures, which will be detected microscopically. After allowing 30 minutes for biodistribution, the animals were anesthetized with a combination of ketamine and xylazine (90:10 mg/kg body weight) via a single intraperitoneal injection. They were then perfused with saline solution and 4% paraformaldehyde to preserve internal organs. The kidneys were then embedded into an approximately 1 cm-diameter 2 cm-long cylindrical agarose block. Optical clearing in agarose was done by first applying dehydration steps with increasing grades of methanol (30 to 100%) to remove water and lipids, then once dehydrated completely, typically after 36-48hours, samples were immersed in BABB (Benzyl Alcohol-Benzyl Benzoate 1:2, Sigma Aldrich) [36] to match refractive index until becoming transparent. Imaging was performed between 1 week and 2 months after preparation.

### Image Acquisition

Following sample preparation, a total of eight healthy and four pathological *ex vivo* kidney scans were acquired using LSFM around eight weeks post-ADPKD induction. Imaging was conducted at the Advanced Digital Microscopy Core Facility (ADMCF) of the Institute for Research in Bioimedicine of Barcelona (IRB Barcelona) with a custom-built LSFM microscope, “MacroSPIM”, previously described and employed for whole organ imaging [38, 39, 40]. In brief, the system enables double sided illumination and double sided detection with two opposing Nikon AZ100 macroscope bodies with Orca Flash 4.0 V3 sCMOS cameras for dual-view imaging, to perform imaging at 4.8x magnification. The excitation light source operated at 561 nm, with emitted fluorescence collected using a BP609/54 filter. Each kidney was scanned by tiled-acquisition, typically 2×3 or 2×4 fields to yield a complete 3D reconstruction of the kidneys. To optimize image quality and reduce scattering degradation, each column was excited with the light sheet that ensured minimal excitation laser propagation through the sample, i.e. right/left column with right or left light sheet respectively. Conversely, each plane was imaged from both sides simultaneously (front and back cameras) to ensure the collection of minimally scattered signals through the cleared kidney by selecting the highest quality image among the two (see next section). The images were acquired with a z-step of 2.5um compared to a lightsheet axial thickness of about 6-7 *µ*m, and the magnification resulted in an XY pixel size of approximately 1.35 *µ*m, compared to the theoretical XY resolution of about 5 to 6µm (estimate in BABB medium, given by the effective numerical aperture of the microscope at 4.8x in air NA=0.153). The final volume was therefore imaged at almost isotropic resolution before subsampling for analysis, and would become isotropic after subsampling. A schematic representation of the whole image acquisition process is shown in Figure 1A.

**Figure 1:**
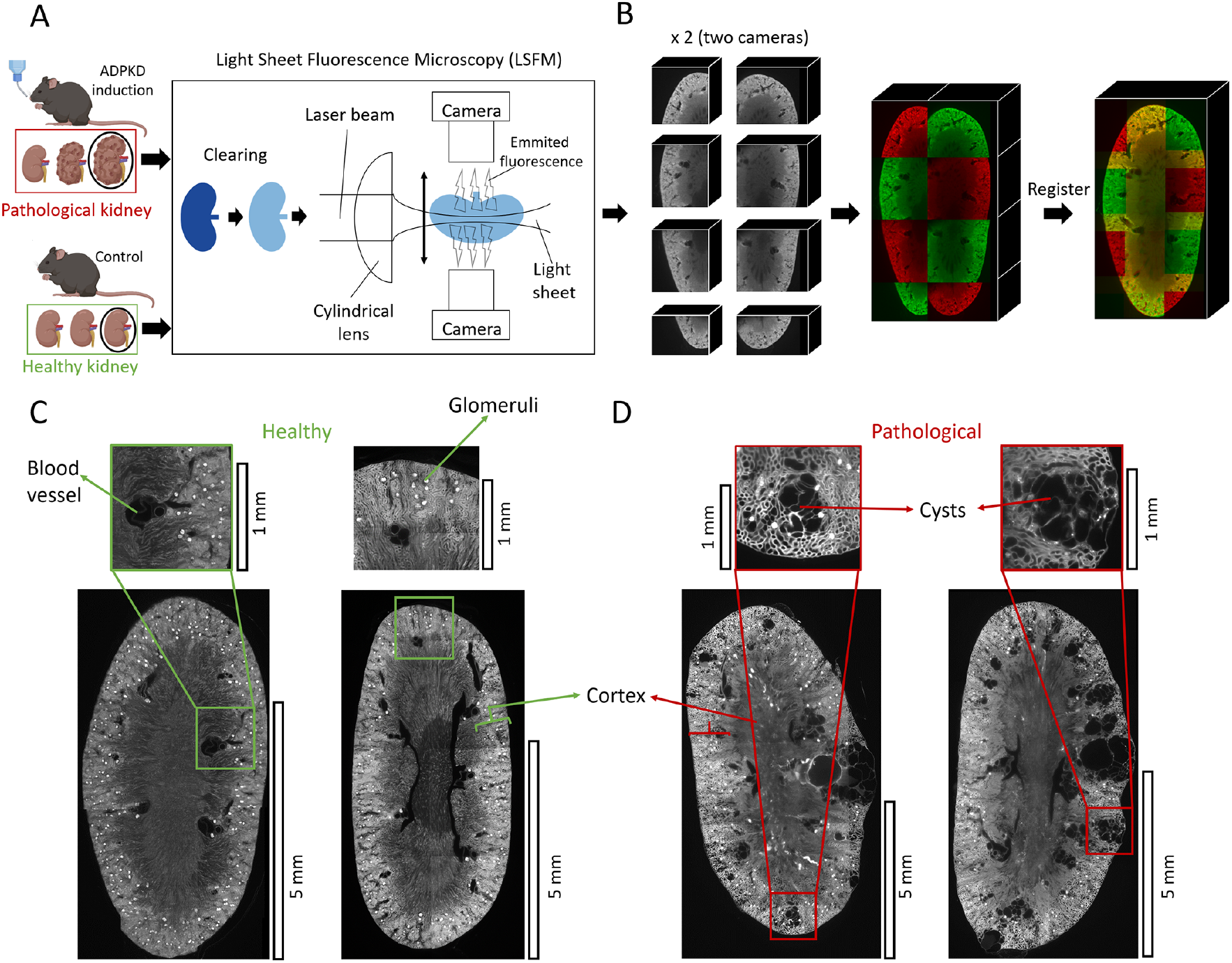
Schematics of the image acquisition workflow and overview of typical LSFM images. (A) ADPKD is induced in a set of mice, with kidney progression illustrated, while control mice are included for comparison. After extraction, kidneys are optically cleared and imaged using a dual-lightsheet, dual-camera MacroSPIM setup, depicted schematically (not at scale). (B) Four sets of overlapping 3D image tiles are acquired per kidney: one for each column with a different lightsheet origin (right or left), repeated for each detection view (0º, 180º, one with each camera). Tile stitching is performed with MosaicExplorerJ [37] by manual positioning of tiles before computing the final 2D stitched planes: first for each column and each camera, then between columns, and finally merging the two opposite views by 2D registration in a selected Z plane. (C-D) LSFM images of healthy (Left, green-annotated) and pathological (Right, red-annotated) kidneys. In healthy tissues, large dark areas represent major blood vessels (see zoomed region at the left). Most glomeruli are concentrated in the outer cortex, which appears brighter in the full kidney image (see second zoomed region highlighting multiple glomeruli and surrounding tubules). In pathological kidneys, ADPKD-induced deformation results in irregular kidney shape and disrupted boundaries due to cyst formation. Cysts appear as abnormal cavities throughout the organ. The first zoomed region (D, left) highlights clustered small cysts distorting the surrounding tissue, with smaller secondary cysts forming nearby. In the second section (D, right), larger cysts visibly disrupt the kidney surface. Scale bars correspond to 5mm on the full kidney images at the bottom panes.

### Image Registration

Individual tiles (datasets approximately 500GB or more) were registered in 2D for each z-plane to generate a single stack of slices for each kidney (half size dataset) to facilitate full-organ analysis. This process was performed using the MosaicExplorerJ [37] script for FIJI [41], specifically designed for handling tiled datasets composed of double illumination and double detection images, including axial correction to compensate for light sheet defects, e.g. axial tilt. Typically, stitching is first optimized by graphically setting tiles to overlap for each column, then each column to one another (e.g. right-to-left stitching), for each camera dataset. Then, front (camera 1) and back (camera 2) image stacks are 2D registered by landmarks selection in a chosen Z-plane (usually the middle plane), hence optimizing image quality by conserving only the proximal images for each camera and discarding the distal images usually more prone to scattering.

The final reconstructed image was downsampled to 50% of its original size in all dimensions for the analysis to reduce computational demands, yielding a new voxel size of 19.659 *µ*m^3^/ voxel. The processed image stacks ranged from 20-35 GB, significantly reducing storage requirements while preserving essential structural details. A schematic representation of this registration process is provided in Figure 1B. In the reconstructed images, cysts appear as distinct structures. Figure 1C shows an example of 2D cross-sections of healthy kidneys, while Figure 1D presents pathological kidneys. Several zoomed-in regions highlight key morphological differences between the tissue types.

### Segmentation of 3D images

After image registration and downsampling, segmentation methods were applied to identify relevant structures and quantitatively assess the effects of ADPKD. The following structures were segmented to characterize disease progression:

#### Full kidney

For each kidney volume, a visual evaluation using FIJI [41] was conducted to determine the most suitable intensity threshold to separate the whole kidney from the background. Due to the high contrast between the organ and the dark background, the image stack was initially thresholded using a global Otsu method, followed by manual adjustments to retain the brightest region and generate an initial full-kidney mask.

A connected component analysis was then performed to preserve only the largest continuous region in each slice, ensuring that only the main kidney body was retained. Morphological operations were subsequently applied to refine the segmentation, specifically a 3D opening and closing operation (using a spherical structural element of 20 voxels radius) to eliminate thin protrusions and small holes. A final 3D closing step (using a larger spherical element of 200 voxels radius) was used to fill larger internal gaps. To optimize computation time, this operation was performed on a 0.25*x* rescaled mask before restoring it to its original size. Padding adjustments were applied to prevent border artifacts.

All morphological operations were executed using Pygorpho [42], a Python library optimized for GPU-based processing of large-scale 3D images. Examples of the resulting segmentation masks are shown in Figure 2A for a healthy kidney and Figure 3A for a pathological kidney, illustrating the full segmented kidney volume.

**Figure 2:**
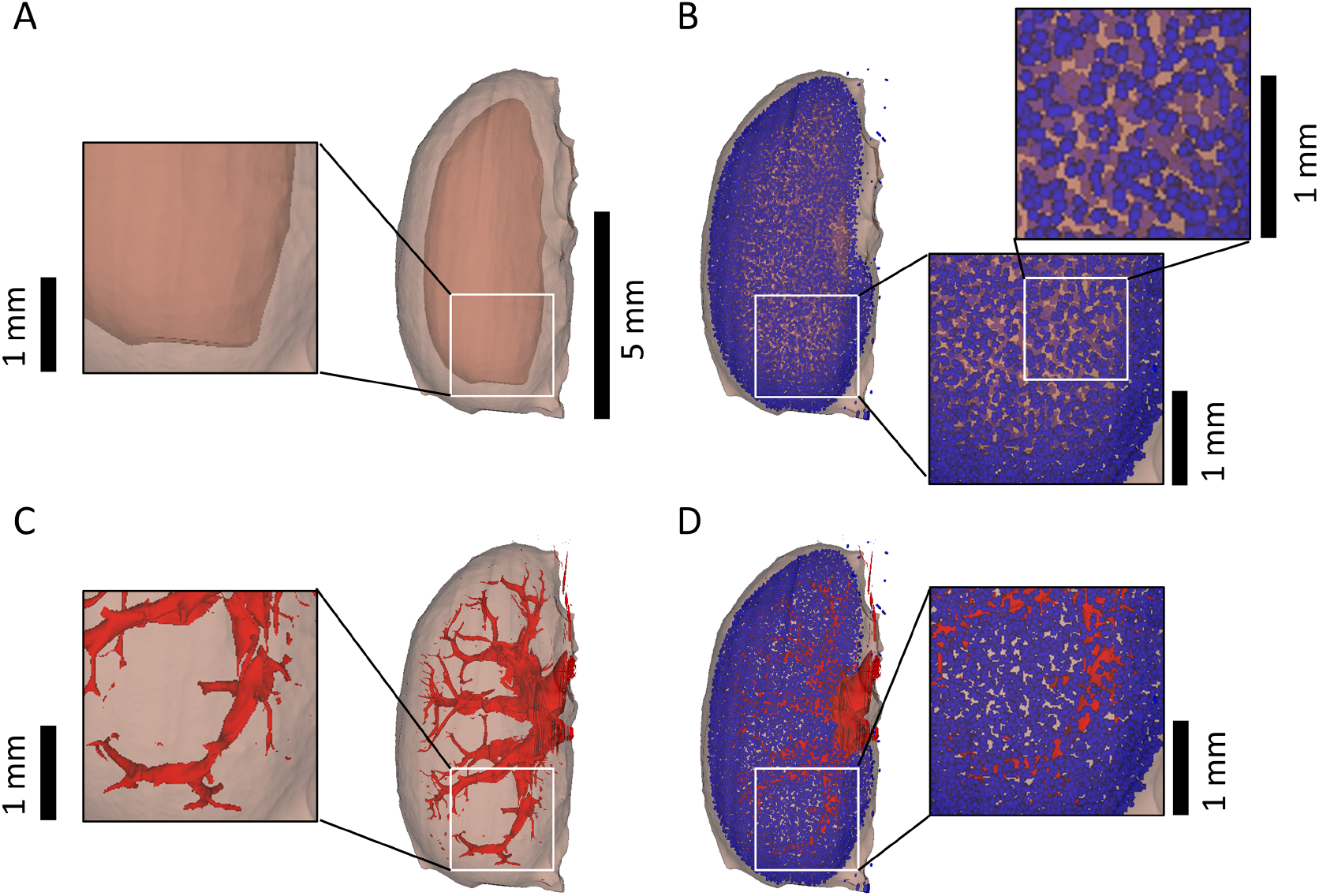
Segmented regions from the LSFM 3D image of a healthy kidney. The entire kidney is shown in brown, with internal structures segmented within it. Zoomed-in regions are provided for illustration. (A) Full kidney segmentation with the cortex in light brown; (B) Glomeruli (blue) overlaid on the cortex (light brown); (C) Internal cavities (red). Main blood vessels appear surrounded by cystic structures, that completely distort the cavities’ structure; (D) Internal cavities (red) and glomeruli (blue). The central scale bar (5 mm) corresponds to the full-organ masks, while the smaller scale bars (1mm) indicate the magnification of the zoomed-in regions.

**Figure 3:**
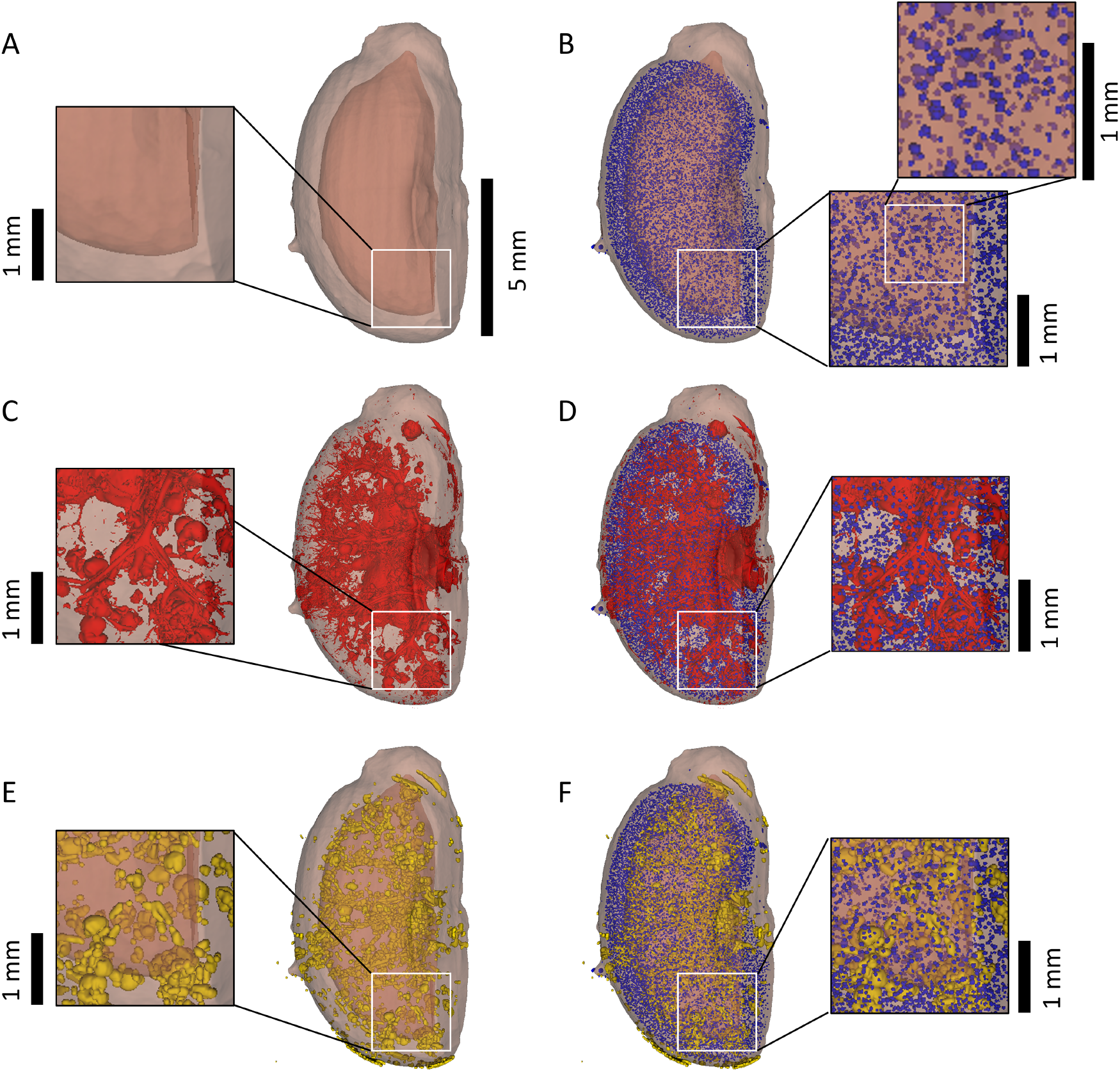
Segmented regions from the LSFM 3D image of a pathological kidney. The entire kidney is shown in brown, with internal structures segmented within it. Zoomed-in regions are provided for illustration. (A) Full kidney segmentation with the cortex in light brown; (B) Glomeruli (blue) overlaid on the cortex (light brown); (C) Internal cavities (red); (D) Internal cavities (red) and glomeruli (blue); (E) Cysts (yellow); (F) Glomeruli (blue) and cysts (yellow). The central scale bar (5 mm) corresponds to the full-organ masks, while the smaller scale bars (1 mm) indicate the magnification for the zoomed-in regions.

#### Cortex

The cortex was approximated as the outer shell of the kidney mask. To delineate this region, the principal axes of each kidney mask were computed. The radius of a spherical structural element was set to half the length of the smallest axis. This structural element was then used to apply 3D morphological erosion to the kidney mask, generating a reduced version representing the inner kidney region. The cortex was approximated as the difference between the original kidney mask and this eroded version.

These cortex masks were generated for both healthy and pathological kidneys, enabling the assessment of structural distributions within the inner and outer kidney regions. Examples of the resulting cortex masks, shown in light brown, can be observed in 2A for a healthy kidney and Figure 3A for a pathological kidney.

#### Internal cavities

In this analysis, the internal cavities were defined as the primary empty spaces within the kidney. In healthy kidneys, this includes the main blood vessel tree, large gaps within the renal pelvis, and other naturally occurring spaces. In pathological kidneys, these structures are present alongside cysts, which appear as empty regions in LSFM images.

Since all internal cavities appeared as dark regions with intensities matching the background, segmentation was performed using Otsu thresholding to distinguish dark areas from brighter structures. The resulting threshold was then manually refined for each kidney to ensure a reliable segmentation of internal cavities while correctly excluding the background. To further refine the mask, the largest connected component in each slice - corresponding to the background - was discarded, preserving only the internal structures of interest.

Examples of the segmented internal cavities are shown in Figure 2C for a healthy kidney, and in 3C for a pathological kidney.

#### Glomeruli

Lectin enabled the visualization of glomeruli as bright, round structures within the kidneys. To accurately detect and quantify these structures, a deep-learning-based segmentation approach was selected. Initially, training a 3D network was considered but ultimately deemed impractical due to the high computational demands and the limited availability of manually annotated 3D training data. Given the distinct circular morphology of glomeruli, a 2D StarDist network [43] was chosen, as it is specifically designed for segmenting round structures, such as cell nuclei. The results of this model differentiate each segmented component by assigning a unique intensity value.

The model was trained on 2D image patches (512 × 512 pixels each). A preprocessing step was introduced to enhance image uniformity. First, a 35-pixel radius mean filter was applied to correct illumination variations. The original image was then divided by the processed version to normalize brightness.

Once trained, the StarDist model was applied to the full dataset, generating segmentation results for individual 2D slices. To construct a 3D glomeruli mask, outputs from multiple slices were combined. An additional post-processing step was implemented to scan the volume slice by slice and identify connected components, after which different glomeruli sections were connected, assigning a unique label to each individual glomerulus across the entire kidney volume.

This segmentation process was applied to all kidney samples. Examples of the glomeruli masks are shown in Figure 2B for a healthy kidney and in Figure 3B for a pathological kidney, with the cortex also depicted. Figures 2D and 3D display the glomeruli masks alongside the segmentation of internal cavities.

#### Cysts

For cyst segmentation, a deep learning approach was also considered to accurately distinguish these structures from surrounding tissue. StarDist was not a suitable choice in this case, as its assumption of convex polygon-shaped elements did not apply - cysts exhibit greater shape variability than glomeruli. Therefore, a U-Net architecture [44] was selected for this task. As with previous segmentations, the network was applied in 2D, with individual slices later combined to generate a 3D cyst mask. The resulting masks were binary, where cysts were labeled as “1” and the background as “0”.

Using the registered image stacks, a series of 2D slices were extracted, and cysts were manually annotated for training and testing. Once trained, the network was applied to pathological kidneys for cyst detection.

To refine segmentation and minimize false positives, several post-processing steps were implemented. First, small misclassified structures were removed using a 2D morphological opening operation with a 10-pixel radius disk. Then, the results were manually reviewed to further improve the cyst masks. After this, the processed slices were iteratively analyzed to label connected components in 3D, ensuring each cyst was assigned a unique intensity value.

Examples of cyst segmentation in a pathological kidney are shown in Figure 3E, while Figure 3F displays the segmented cysts alongside the glomeruli.

### Measurements from the Segmentation Masks

Following segmentation, a series of quantitative measurements were extracted to analyze kidney structure and pathology. for each full kidney mask, the centroid coordinates, total volume, and eigenvalues of the inertia tensor were obtained. These eigenvalues were used to calculate the lengths of the three principal axes by approximating the kidney as a 3D ellipsoid, what would be used for the cortex segmentation.

For the internal cavities masks, the total volume of the segmented cavities was computed, offering a quantitative assessment of empty spaces within the kidney. In the glomeruli segmentations, the centroid coordinates and individual volumes of each glomerulus were recorded, enabling a detailed evaluation of their distribution and size. Similarly, for the cyst segmentations, the centroid coordinates and volumes of each cyst were measured, facilitating the characterization of cystic expansion in pathological kidneys.

### Texture analysis

Besides the segmentations obtained from the images, a texture analysis was conducted. Patches of 128×128×128 voxels were extracted from cubic regions fully contained within the cortex of the 3D LSFM full kidney images (Figure 4). Each patch was randomly centered in one of the previously segmented glomeruli to facilitate direct comparisons between them. In pathological kidneys, an additional constraint was applied: the path center was required to be within a radius of 128 voxels of a cyst centroid. This ensured that the selected region to be in close proximity to fully developed cysts, where tubular deformation is most evident. The extracted patches were verified to ensure they accurately represented tubules in their respective regions.

**Figure 4:**
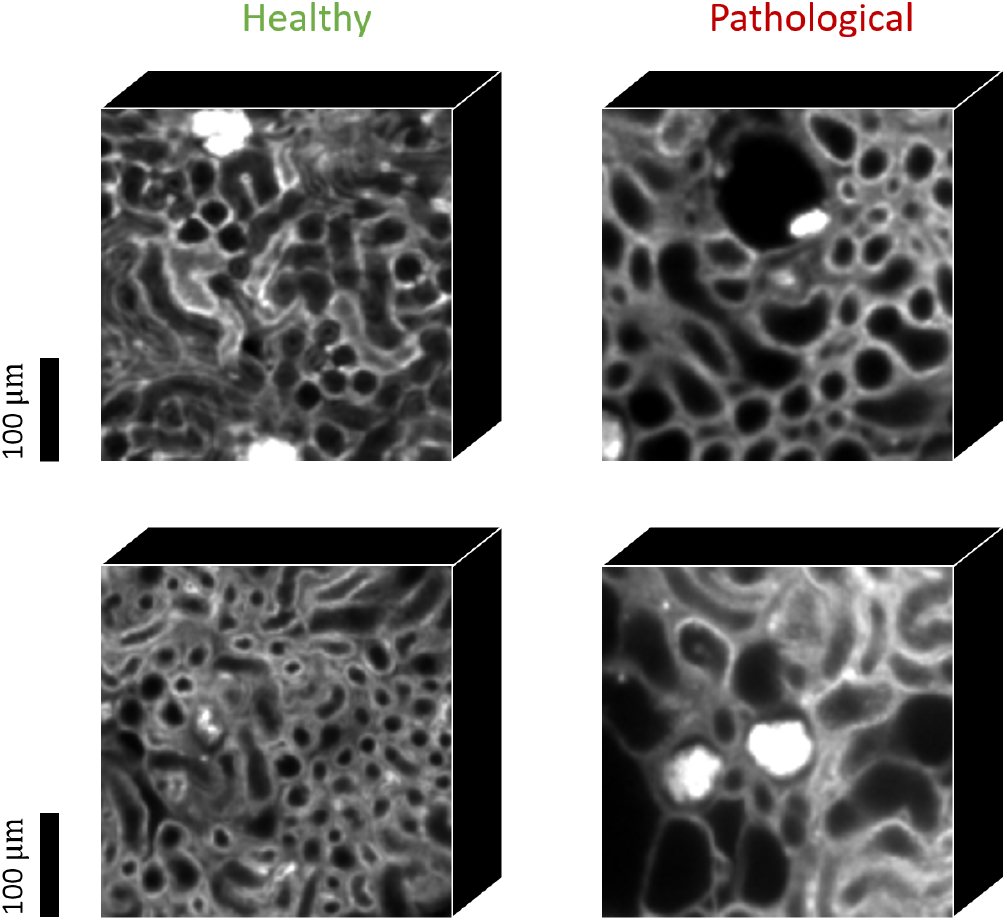
Examples of healthy and pathological patches from the cortex for texture analysis. 2D slices from 3D 128×128×128 patches, surrounding cysts in the case of pathological patches. Healthy patches show smaller tubules with more ordered structures, while the apparition of cysts on pathological patches causes surrounding tubules to increase in size, being deformed in the process. Scale bar: 100 µm.

To standardize intensity variation, each patch was normalized to a [0, 1] intensity range, and a set of texture features was extracted. These features were concatenated into a single feature vector, serving as a compact representation of the image for classification. The selection of 12 texture features aimed to provide a comprehensive yet efficient characterization, balancing representational power with dimensionality constraints to prevent excessive feature complexity.

The first four extracted features captured fundamental intensity-based properties:

1. **Mean Intensity:** Represents the average pixel intensity in the image, providing a measure of overall brightness (values ∈ [0,1]).
2. **Intensity Variance:** Measures the spread of pixel intensities around the mean, indicating overall contrast (values ∈ [0,0.25]).
3. **Intensity Skewness:** Quantifies the asymmetry of the pixel intensity distribution (any value).
4. **Intensity Kurtosis:** Describes the “tailedness” of the intensity distribution, where higher values indicate a sharper peak and heavier tails (values ≥ 1).

Four wavelet features were extracted for edge detection:

5. **Wavelet Approximation Coefficient Mean:** Computed from the approximation coefficients of a 3D discrete wavelet transform, this feature represents the average value of the low-frequency components, capturing the overall structure of the image (values ∈ [0,1]).
6. **Wavelet Horizontal Detail Coefficient Mean:** Represents the mean value of the horizontal detail coefficients from the 3D wavelet transform, providing information about edge details in the horizontal direction (values ∈ [-1,1]).
7. **Wavelet Vertical Detail Coefficient Mean:** Captures the edge information in the vertical direction (values ∈ [-1,1]).
8. **Wavelet Depth Detail Coefficient Mean:** Captures edge information in the depth direction (values ∈ [-1,1]).

Two Fourier domain features for frequency analysis:

9. **Fourier Dominant Frequency Magnitude:** Derived from the 3D Fast Fourier Transform (FFT), this feature represents the magnitude of the dominant frequency, indicating the most significant repeating pattern in the image (values ≥ 0).
10. **Total Energy in Frequency Domain:** Computed as the sum of squared magnitudes of the FFT coefficients, this feature quantifies the total energy of spatial variations in the image (values ≥ 0).

Two additional features offering more information on local patterns and cavity proportion for assessing tissue heterogeneity:

11. **Local Binary Pattern (LBP) Uniformity:** Measures the uniformity of local binary patterns in a 3D image, calculated from the histogram of LBP values across the image. Higher uniformity indicates more homogeneous texture patterns (values ∈ [0,1]).
12. **Porosity:** Defined as the ratio of low-intensity pixels (holes) to the total number of pixels in the image [45]. It is computed by thresholding the image using the mean intensity and dividing the count of low-intensity pixels by the total pixel count (values ∈ [0,1]).

Following feature extraction, an XGBooster classifier [46] was trained using the generated feature vectors to classify patches as healthy or pathological. The classification process also provided insight into the relative importance of each feature, helping to identify key indicators of the disease.

To optimize performance, a hyperparameter grid search was conducted to determine the best model configuration. The classifier was trained on a fixed training set, testing different hyperparameter combinations, and evaluated on a constant validation set. The configuration yielding the highest validation score after 100 training epochs was then applied to a separate test set for the final classification performance assessment.

The texture features used for classification were ranked based on their Gain-based importance, which quantifies the contribution of each feature to the predictive performance of the model. This metric is computed by summing the gain values from all decision tree nodes where a given feature is used to split the data. A higher importance score indicates a greater contribution to the predictive power of the model, meaning that variations in this feature play a more significant role in distinguishing between pathological and healthy tissue. The importance scores are normalized, with their sum equaling one.

### Statistical Methods

All measurement sets obtained from the segmentation masks were positively tested for normality using the Shapiro-Wilk test at a 95% confidence level. For full kidney volume, a single value was extracted per kidney, and the mean and standard deviation were computed separately for healthy and pathological specimens. To assess significant differences between groups, a t-test for the difference of means at 95% confidence was conducted. Cavity volume proportions were calculated by dividing the cavity volume by the total kidney volume, yielding a single proportion per organ. The mean and standard deviation of these proportions were computed for both groups, followed by the same t-test for statistical comparison.

The number of glomeruli was measured for each kidney, and the mean and standard deviation were computed separately for healthy and pathological groups. A t-test at 95% confidence was applied to compare the means.

For glomerular spatial distribution, the centroid coordinates of all glomeruli within each kidney were used to calculate the nearest-neighbor distance for each glomerulus. The mean and standard deviation of these distances were then computed within each kidney. Two separate analyses were conducted:

1. **Inter-kidney analysis**: The mean nearest-neighbor distance per kidney was used to compute the group mean and standard deviation across healthy and pathological specimens, followed by a t-test to compare the groups.
2. **Intra-kidney variability analysis**: The standard deviation of nearest-neighbor distances per kidney was collected across specimens, and a group-level mean, standard deviation, and t-test were computed.

The proportion of glomeruli in the cortex was determined by counting the number of glomerular centroids within the cortical region and dividing it by the total glomerulus count for each kidney. These proportions were aggregated to compute the mean and standard deviation for healthy and pathological groups, with a t-test applied for comparison.

Similarly, the proportion of cysts in the cortex was calculated by dividing the number of cyst centroids within the cortex by the total cyst count in each kidney. Finally, the mean cyst volume per kidney was obtained by averaging the volumes of all cysts within a given kidney. These mean cyst volumes were then aggregated across specimens to compute the overall mean and standard deviation.

P-values of the t-tests can be observed at Figure 5.

**Figure 5:**
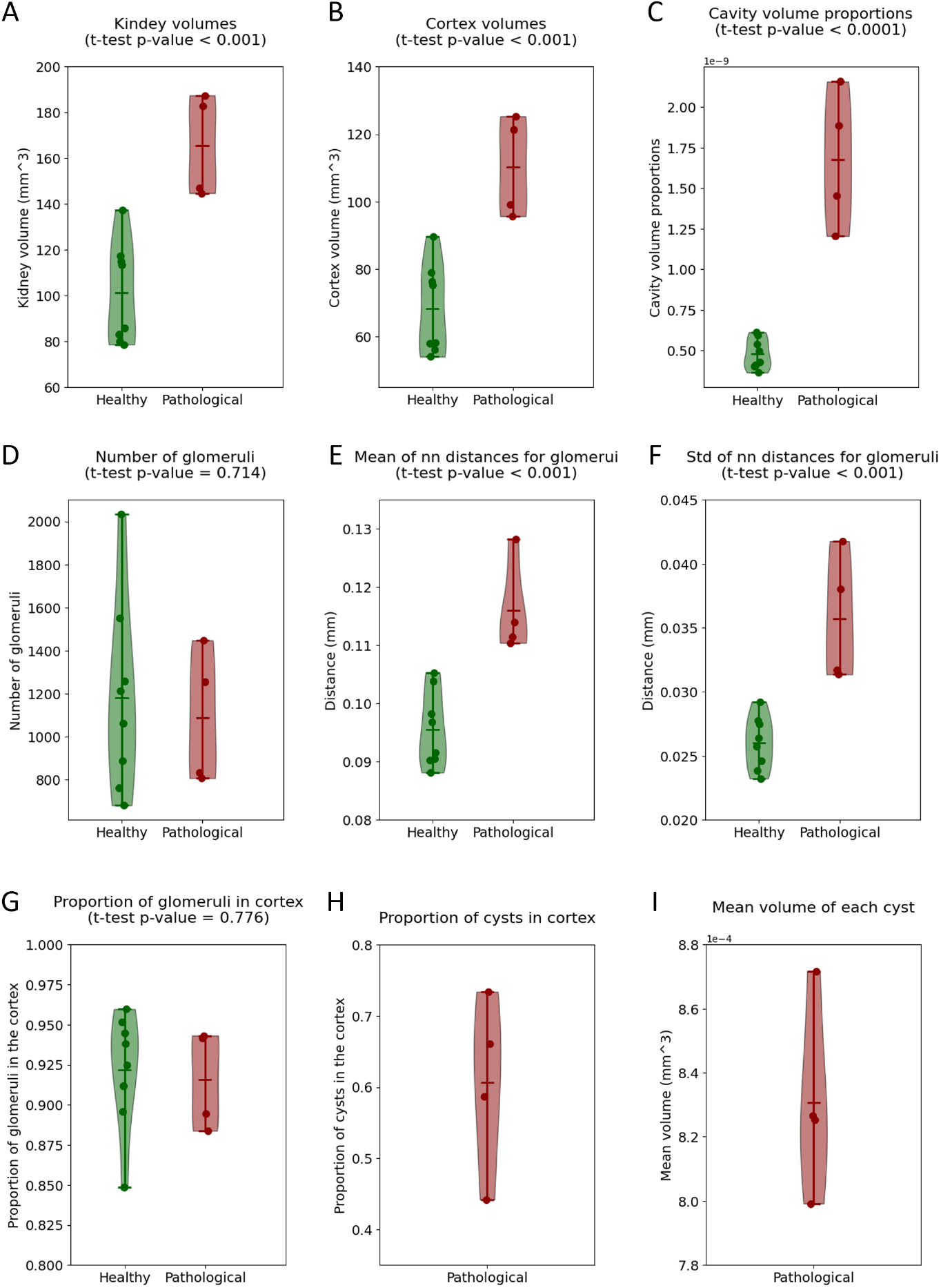
Violin plots of different measurements taken from the segmentation of LSFM kidney images. Measures from healthy kidneys are shown in green, and from pathological in red. The central horizontal bar of each group represents the median, and individual data points are shown as dots. In the cases where a t-test for the difference in means was performed between healthy and pathological, the p-value is shown in the title. A. Kidney volumes; B. Cortex volumes; C. Cavity volume proportions. D. Number of glomeruli within each kidney; E. Mean distances to the glomeruli nearest neighbor for each kidney; F. Standard deviations of the distance to the glomeruli nearest neighbor for each kidney. G. Proportion of glomeruli in the cortex; H. Proportion of cysts in the cortex I. Mean volume for each individual cyst.

### Hardware

All computations were performed using Python on a computer with 128GB RAM and an RTX3090 GPU.

## 3 Results

### Segmentation Measurements

Quantitative measurements were extracted from the segmentation masks to provide a detailed analysis of kidney morphology. Several parameters were computed for both healthy and pathological groups, with p-values from statistical tests displayed in the titles of the respective graphs.

Figure 5A presents the total kidney volume distribution for both groups. The healthy kidneys exhibit a significantly smaller size, with a mean volume of 100.34 ± 20.48 mm^3^, compared to the pathological kidneys, which have a mean volume of 163.86 ± 20.47 mm^3^. The low p-value from the t-test confirms a significant difference between the two groups. On the other hand, defining the exact boundary of the cortex is challenging due to its gradual transition into deeper kidney structures. In this study, it was approximated as the outer shell of the kidney mask. Figure 5B illustrates the cortex volume distribution. A significant difference is also observed, with the healthy group showing a mean cortex volume of 68.33 ± 12.45 mm^3^, while the pathological group exhibits an increased mean volume of 110.34 ± 13.11 mm^3^. The results clearly demonstrate an increase in overall kidney volume in pathological kidneys.

Similarly, cortex volume is also noticeably larger in pathological kidneys, correlating directly with the increase in total kidney volume. These changes indicate that cyst formation and expansion enlarge the kidney, displacing pre-existing structures.

Figure 5C displays the cavity volume proportion. Although the values are small for both groups, they are notably lower in healthy kidneys (4.75 × 10^*−*10^ ± 8.71 × 10^*−*11^), while the pathological group shows a significant increase (1.65 × 10^*−*9^ ± 3.66 × 10^*−*10^), as confirmed by the t-test results. This measure highlights how ADPKD progressively transforms a normal kidney - characterized by a functional vascular system and natural tissue gaps - into a highly cystic organ with a substantial increase in internal cavity space.

Another key structure analyzed in this study is the glomeruli. For their segmentation, a StarDist model was trained using 2D patches extracted from the images, with manual annotation used to construct a ground truth (GT) dataset. A total of 198 patches were used for training and 22 for validation over 400 epochs, utilizing pretrained weights from the DSB 2018 model. The network was then tested on 45 patches, achieving a mean F1-score of 0.93 ± 0.07.

After segmenting the glomeruli using the trained network, the total number of glomeruli per kidney was computed and is shown in Figure 5D. The mean number of glomeruli in healthy kidneys was 1181.12 ± 417.25, while in pathological kidneys, it was 1086.00 ± 274.68. In this case, no significant difference was observed between the two groups.

For this analysis, nearest-neighbor distances were computed for all glomeruli within each kidney, and a single representative value per kidney was obtained by averaging these distances. These values were then aggregated across all kidneys for final statistical analysis. This method was preferred over directly pooling all glomeruli across kidneys, as each kidney exhibits distinct size and morphological differences. Treating each kidney as an independent unit allowed for more accurate characterization of individual organ structures, preserving the biological independence of each specimen. Figure 5E presents the mean nearest-neighbor distance for each kidney, which quantifies how closely glomeruli are clustered. Unlike the total glomerular count, there was a clear increase in nearest-neighbor distance in pathological kidneys (0.116 ± 0.007 mm) compared to healthy specimens (0.096 ± 0.006 mm), with a statistically significant p-value. Additionally, Figure 5F displays the standard deviation of nearest-neighbor distances, providing insight into heterogeneity in glomerular distribution. Pathological kidneys exhibited a higher mean standard deviation (0.036 ± 0.004 mm) compared to healthy kidneys (0.026 ± 0.002), with this difference also being statistically significant.

The total number of glomeruli remains relatively constant between healthy and pathological kidneys, suggesting that ADPKD does not directly cause glomerular loss. However, the mean nearest-neighbor distance between glomeruli is significantly larger in pathological kidneys, indicating a more dispersed glomerular distribution. This suggests that as cysts form and expand, the surrounding tissue undergoes structural deformation, pushing glomeruli apart. The mean nearest-neighbor distance between glomeruli serves as a quantitative indicator of structural deformation, which is expected to worsen as the disease progresses. Additionally, the standard deviation of the nearest neighbor distance is significantly larger in the pathological specimens, reinforcing the observation that glomerular distribution becomes increasingly heterogeneous due to ADPKD-driven tissue remodeling.

For cyst segmentation, a U-Net model was trained using 43 masks for training and 26 for validation over 100 epochs. Additional training beyond this point did not improve performance. The model was tested on 12 images, achieving a mean Intersection over Union (IoU) of 0.71 ± 0.07.

To evaluate the spatial distribution of cysts and glomeruli, the proportion of these structures within the cortex was analyzed. For glomeruli, Figure 5G compares de results across groups, showing that the proportion of glomeruli within the cortex was nearly identical in both groups (0.92 ± 0.03 in both healthy and ADPKD-induced kidneys), indicating that the majority of glomeruli were located within the cortex for both healthy and pathological specimens. Visual inspection of the LSFM images clearly shows that these spheroids are primarily situated in the outer region, with their corresponding tubules extending inward. No significant difference in glomerular distribution was observed between the two groups, suggesting that ADPKD does not induce sufficient deformation to alter their average location. Although glomeruli are pushed farther apart due to tissue expansion from cyst formation, they remain predominantly in the cortex throughout disease progression.

In contrast, the proportion of cysts within the cortex among pathological kidneys exhibited high variability (5H), with a mean proportion of 0.61 ± 0.11. This indicates that cyst formation does not seem to be strongly related to the depth within the kidney. Cysts appear throughout the organ, but a mean proportion exceeding 0.5 suggests a slight tendency for cysts to form more frequently in the cortex.

Additionally, Figure 5I presents the mean volume of individual cysts. To maintain organ-specific individuality, the mean cyst volume was first computed per kidney and then aggregated across specimens. The results show a mean of 8.2 × 10^*−*4^ ± 2.58 × 10^*−*5^ mm^3^ in pathological kidneys, though some kidneys contained noticeably larger cysts than others.

### Texture Analysis

After visually inspecting the 3D full kidney images, it could be noticed that ADPKD produced some modifications in the dense network of tubules that become slowly deformed as the disease progresses, increasing their thickness and eventually leading to cyst apparition. Segmentation of the complete tubular network was out of the scope of the project, so to characterize these changes, a texture analysis on these areas was conducted as a complement to the previous segmentations. This was performed on 3D patches extracted from the cortex area of the full organ images, where the changes in tubule structures can be better observed. Using a series of healthy as well as pathological patches, we aimed to assess the importance ranking of different texture features on the characterization of cortex tissue as pathological. This analysis serves as a basis for further evaluation of the evolution of ADPKD in the tissue over time, as well as the effects of low-scale treatments. An example of some of these patches, both healthy and pathological, can be observed in Figure 4, where the deformations produced by cyst formation on tubules are visible.

A total of 477 patches were used for this analysis. After extracting the corresponding feature vectors, the dataset was divided as follows: 317 vectors were used in the training set (237 healthy, 80 pathological), 80 for validation (40 healthy, 40 pathological), and 80 for testing (40 healthy, 40 pathological).

The optimal hyperparameter values obtained from the XGBoost grid search are highlighted in bold in Table 1. When applying the best-trained model to the test set, it achieved an F1-score of 0.90, with a precision of 0.92 and a recall of 0.88 (considering pathological as the positive class). These results proved its ability to effectively differentiate pathological from healthy kidneys based on the extracted texture descriptor, which was deemed sufficient to justify the use of this classifier to evaluate the relative importance of individual texture features in the classification process.

**Table 1:** Hyperparameter Grid Search Values for XGBoost Classifier. Optimal values after tuning highlighted in Bold.

| Hyperparameter | Values |
| --- | --- |
| Number of Estimators | 50, **100**, 200 |
| Maximum Depth | 3, 6, **9** |
| Learning Rate | 0.01, **0.1**, 0.2 |
| Subsample | 0.6, 0.8, **1.0** |
| Column Subsample by Tree | 0.6, 0.8, **1.0** |
| Minimum Loss Reduction | **0**, 0.1, 0.2 |
| Minimum Child Weight | **1**, 3, 5 |
| L1 Regularization Term | **0**, 0.01, 0.1 |
| L2 Regularization Term | 1, 1.5, **2** |

Figure 6A presents the feature importance scores obtained from the classifier. The results indicate that LBP uniformity was the most relevant feature, followed by porosity and mean intensity, while the remaining features exhibited lower importance scores. The Wavelet Approximation Coefficient Mean and Fourier Dominant Frequency Magnitude contributed the least, with no measurable impact on classification.

**Figure 6:**
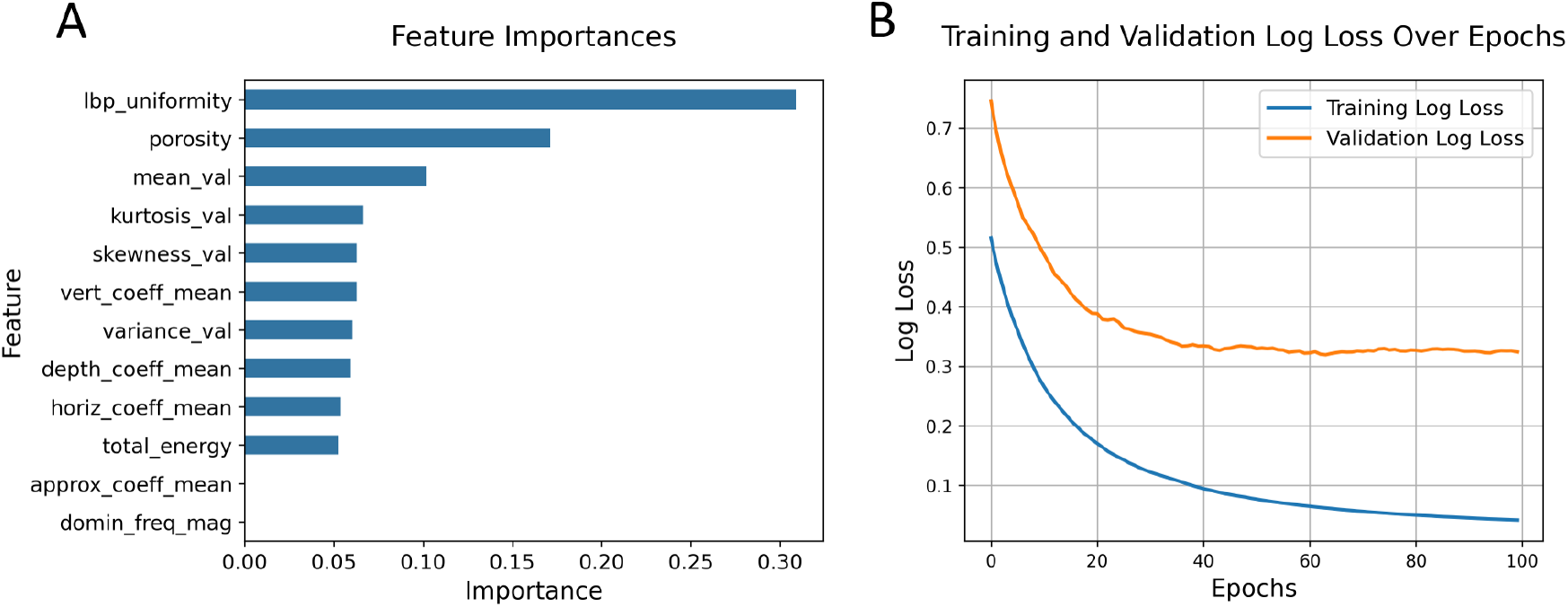
Texture analysis results. (A) Ranking of texture features based on their contribution to the classification model, in descending order of importance: LBP Uniformity (lbp uniformity), Porosity, Mean Intensity (mean val), Intensity Kurtosis (kurtosis val), Intensity Skewness (skewness val), Wavelet Vertical Detail Coefficient Mean (vert coeff mean), Intensity Variance (variance val), Wavelet Depth Detail Coefficient Mean (depth coeff mean), Wavelet Horizontal Detail Coefficient Mean (horiz coeff mean), Total Energy in Frequency Domain (total energy), Wavelet Approximation Coefficient Mean (approx coeff mean) and Fourier Dominant Frequency Magnitude (domin freq mag). (B) Log-loss evolution during training of the best-performing XGBoost model for the training and validation sets.

Figure 6B illustrates the log-loss evolution for both training and validation sets during the training of the best-performing XGBoost model.

## 4 Discussion

This manuscript presents a novel application of LSFM for the evaluation of ADPKD effects on mouse kidneys. In recent years, LSFM has been used sporadically to analyze the full kidney. For instance, [32] utilized automated image analysis to assess glomerular hypertrophy in a mouse model of diabetic nephropathy, which shows similarities to the current analysis, while [31] also segmented glomeruli in nephritic kidneys. Our processing pipeline differs from them in its integration with deep learning as an alternative for glomeruli segmentation, as well as cysts.

In terms of the study of ADPKD, this analysis presents an alternative to more commonly used imaging methods at several levels. The most commonly used measure of ADPKD evolution is Total Kidney Volume (TKV). While normally calculated from MRI images [5], we could extract this parameter from our LSFM reconstructions, showing the suitability of the method to observe the disease’s development through this whole-organ feature. On the other hand, in terms of ADPKD animal studies, histology imaging is currently the norm, allowing to extract cystic index measurements to observe the growth of cysts through the organ, as shown in studies like [14] and [15]. On that note, the cavity volume calculated from LSFM images can be seen as a 3D alternative to the 2D cystic index measurements taken in histology imaging studies. As the disease advances, these cavities replace normal tissue, serving as a useful metric to assess the extent of kidney degradation. Histological studies at later stages of the disease reveal that cyst proliferation eventually saturates the entire kidney, leaving minimal intact tissue [18]. LSFM imaging of later-stage kidneys is expected to yield similar three-dimensional reconstructions. However, preparing samples from late-stage kidneys may present challenges, especially if the structural integrity of the tissue is compromised. The texture analysis of tubular structures offers additional insight into the structure of the organ at a lower level than that of previous volumetric techniques on ADPKD studies, allowing us to better observe the specific effects produced by this condition. By identifying several texture descriptors that explain the changes produced in the tubular structure, we present a way to evaluate the evolution of ADPKD-induced morphological changes in the compact structure of the kidney. Namely, LBP uniformity and porosity, as well as similarly defined metrics, can serve as markers of the disease progression moving forward to more detailed low-level studies.

What is more, in addition to quantitative measurements that help assess the effects of ADPKD at a finer scale, LSFM has been shown to provide a high-resolution informative visualization of kidney architecture. The ability to visually navigate through different 3D structures of this organ in different conditions provides a clear representation of both micro- and macroscopic changes induced by the disease, serving as well as a powerful tool for understanding ADPKD progression.

In this study, we used fluorescent-lectin-mediated labeling to reveal the blood vessels, and we analyzed a single fluorescence channel. The lectin labeling was efficient in revealing glomeruli, but the final fluorescence intensity of blood vessels was somehow weak, compared to the co-existing autofluorescence, to enable reliable segmentation of the finer vessels/capillaries for segmentation. This enabled to use the same fluorescence channel for both segmentation and texture analysis, thereby exploiting autofluorescence that contributes positively to the texture analysis. However, the use of different clearing protocols, in particular those that enable to remove autofluorescence, could yield more specific data for the purpose of blood vessel segmentation. Finally, acquisitions were performed at moderate magnification (4.8x), to provide contained image data size. Imaging at the maximum available resolution of the instrument, e.g. at higher magnification (up to 24x) with a thinner lightsheet (down to 3-4 *µ*m instead of about 6-7 *µ*m here), would have yielded larger data volumes (about 50 times larger volumes, over 10TB per sample), hence has not been prioritized. Nevertheless, the large size of LSFM images posed a significant computational challenge, even after downsampling to 50% of their original size. While GPU acceleration was utilized for deep learning training and morphological operations, implementing a high-performance computing (HPC) workflow would further optimize full-resolution image processing. More detailed information on specific kidney regions could potentially be extracted directly from the original-size image tiles. This would require the development of a pipeline capable of handling full-resolution data without excessive memory constraints. Rather than processing the entire volume at once, a bricking approach or distributed computation would enable localized analyses while maintaining data integrity. Implementing such strategies on a more powerful computational platform would facilitate the processing of both individual tiles and full datasets, ensuring detailed structural insights without performance bottlenecks.

A limitation of this study was the small number of samples, primarily determined by specimen availability. Future studies should aim to increase this amount to strengthen statistical conclusions and further validate the findings. Expanding the dataset to include kidneys at different stages of ADPKD progression would also provide a more comprehensive understanding of disease evolution and improve statistical robustness, allowing as well to assess treatment efficacy in preclinical models over time.

In summary, this work highlights the effectiveness of LSFM in producing high-resolution, three-dimensional reconstructions of mouse kidneys, offering a comprehensive view of tissue deformation and cyst formation in ADPKD. LSFM enables detailed comparisons between healthy and pathological kidneys, revealing significant differences in organ volume, internal cavity proportions, and glomerular distribution.

Key findings include:

- A marked increase in kidney volume and internal cavities in ADKD-affected kidneys, primarily driven by cyst formation.
- Glomerular count remains relatively stable, but glomeruli become more dispersed due to structural deformation.
Cyst formation is slightly more frequent in the cortex, though cysts appear throughout the kidney.
- Texture analysis of small 3D patches identified LBP uniformity and porosity as key indicators of low-scale ADPKD progression in LSFM images.

These results suggest that texture-based analyses, particularly those evaluation pattern uniformity and structural porosity, could be valuable for characterizing ADPKD progression in future longitudinal studies using LSFM imaging.

## 5 Conclusions

LSFM has proven to be a valuable tool for studying the structural changes caused by ADPKD in mouse kidneys. With its high-resolution capabilities, LSFM enables the acquisition of measurements comparable to those obtained from high-quality 3D histology images while offering a more comprehensive representation of the organ’s true condition. Unlike traditional histology, which provides information from isolated bidimensional sections, LSFM allows for the analysis of entire volumetric structures, preserving spatial relationships within the kidney.

This technique also provides more detailed visualization than MRI or CT, particularly for small organs, making it ideal for low-level studies of fine structures such as tubules. Additionally, LSFM can be used to track fluorescently labeled particles within the kidney, expanding its applications in drug delivery and disease modeling.

In this study, the quantification of ADPK-induced structural changes was successfully performed through segmentation and texture analysis, leveraging machine learning techniques. These results serve as a foundation for future evaluations of ADPKD treatments.

Moving forward, future research should focus on:

- Expanding sample sizes to enhance the statistical robustness of findings.
- Conducting longitudinal studies to track disease progression over time.
- Exploring translational applications, particularly in human kidney disease models.
- Monitoring targeted drug distribution, including Adeno-Associated Viruses (AAVs) for gen therapy, using LSFM capability to detect fluorescently labeled therapeutic agents.

With these advancements, LSFM could become a key imaging tool for both preclinical and translational research, facilitating new insights into ADPKD progression and treatment responses. The ability to visualize and quantify disease-related structural change at high resolution positions LSFM as a promising technique for evaluating novel interventions, such as targeted drug delivery or gene therapy strategies, ultimately contributing to improved nephrology research and patient outcomes.

## 6 Acknowledgements

This work was partially funded by Ministerio de Ciencia, Innovación y Universidades, through an FPU fellowship (FPU19/02854). Additional funding was provided by Ministerio de Ciencia, Innovación y Universidades, Agencia Estatal de Investigación (MCIN/AEI/10.13039/501100011033/), under Grants PID2021-128862OB-I00, PID2023-152631OB-I00, TED2021-129392B-I00 and RTC-2017-6600-1, co-financed by European Regional Development Fund (ERDF), “A way of making Europe”. Partial support also came from Grant 0011-1411-2019-000074 from proyectos de I+D estratégicos (RIS3) from Departamento de Desarrollo Económico del Gobierno de Navarra. Part of the funding was also provided by CIBER de Salud Mental - ISCIII (project number CB07/09/0031); Delegación del Gobierno para el Plan Nacional sobre Drogas correspondiente a fondos del Mecanismo de Recuperación, Transformación y Resiliencia de la Unión Europea (EXP2022/008917); and Delegación del Gobierno para el Plan Nacional sobre Drogas (2024I090) The authors used ChatGPT (OpenAI) for language refinement and editorial assistance in drafting sections of the manuscript. All intellectual contributions, scientific content, and conclusions were formulated by the authors. All co-authors reviewed and approved the final manuscript to ensure accuracy and originality.

## 7 Ethics Approval

Experimental procedures conducted during this project adhered to the European Communities Council Directive (2010/63/EU) and national rules (RD53/2013, ECC/566/2015) for the care of laboratory animals. All of the procedures involving mice have been approved by the Universidad de Navarra Animal Research Ethics Committee under the animal protocol number 081c-19.

## 8 Data sharing

The code used during this study is available at https://github.com/pdelgado248/LSFM-for-ADPKD. Due to the very large size of the LSFM images used during the project, they are available upon request from the authors.

## 9 Author Contributions

PDR wrote the manuscript, annotated data, performed segmentations and further data analyses, as well as the texture analysis; IV annotated data, registered the images, and contributed to the segmentations; GRM contributed in the texture analysis methods; LB performed samples clearing; NLR and MLSM provided the samples and annotated data; LNS annotated data; JS contributed to the development of segmentation methods; RA developed the animal model and provided the specimens; JC developed the light sheet instrument and designed the imaging workflow, acquired the images, and provided insight on the analyses to be performed; AMB annotated data, coordinated the project; provided the computational resources. All authors reviewed the manuscript.

